# A novel ultra high-throughput 16S rRNA amplicon sequencing library preparation method on the Illumina HiSeq platform

**DOI:** 10.1101/124057

**Authors:** Eric J. de Muinck, Pål Trosvik, Gregor D. Gilfillan, Arvind Y. M. Sundaram

**Affiliations:** Centre for Ecological and Evolutionary Synthesis, Dept. of Biosciences, University of Oslo, Oslo, Norway; Department of Medical Genetics, Oslo University Hospital and University of Oslo, Oslo, Norway

**Keywords:** 16S amplicon sequencing, Illumina library preparation, indexed PCR, mock community, environmental sequencing, benchmarking, PCR bias, chimera formation

## Abstract

**Background:** Advances in sequencing technologies and bioinformatics have made the analysis of microbial communities almost routine. Nonetheless, the need remains to improve on the techniques used for gathering such data, including increasing throughput while lowering cost, and benchmarking the techniques so that potential sources of bias can be better characterized.

**Results:** We present a triple-index amplicon sequencing strategy that uses a two-stage PCR protocol. The strategy was extensively benchmarked through analysis of a mock community in order to assess biases introduced by sample indexing, number of PCR cycles, and template concentration. We further evaluated the method through re-sequencing of a standardized environmental sample. Finally, we evaluated our protocol on a set of fecal samples from a small cohort of healthy adults, demonstrating good performance in a realistic experimental setting. Between-sample variation was mainly related to batch effects, such as DNA extraction, while sample indexing was also a significant source of bias. PCR cycle number strongly influenced chimera formation and affected relative abundance estimates of species with high GC content. Libraries were sequenced using the Illumina HiSeq and MiSeq platforms to demonstrate that this protocol is highly scalable to sequence thousands of samples at a very low cost.

**Conclusions:** Here, we provide the most comprehensive study of performance and bias inherent to a 16S rRNA gene amplicon sequencing method to date. Triple-indexing greatly reduces the number of long custom DNA oligos required for library preparation, while the inclusion of variable length heterogeneity spacers minimizes the need for PhiX spike-in. This design results in a significant cost reduction of highly multiplexed amplicon sequencing. The biases we characterize highlight the need for highly standardized protocols. Reassuringly, we find that the biological signal is a far stronger structuring factor than the various sources of bias.

## Background

Significant advances in microbiome analysis have been achieved by the adoption of improved sequencing technologies and increased computational capabilities. Sequencing large numbers of relatively short DNA fragments has become routine, and microbiologists have adapted these technologies to characterize communities of microbes by either targeted sequencing of conserved regions containing phylogenetically informative polymorphisms (e.g. 16S or 18S rRNA gene sequencing), or by sequencing a sub-set of the randomly sheared DNA molecules in a sample (so-called shotgun metagenomics). Both approaches present unique challenges for identification and interpretation of biologically meaningful information, and for the moment, the high costs associated with deep sequencing in shotgun metagenomics currently limits full exploitation. As such, combined approaches may offer the best possibility for achieving a significantly improved understanding of complex microbial communities. Thus, targeted PCR amplification and sequencing of the 16S rRNA gene continues to offer a powerful and economic way to gain insight into the bacterial community composition in large numbers of samples.

To address issues related to accuracy, cost, and throughput, we have designed an improved Illumina compatible library preparation protocol, combining features of several existing popular sequencing strategies, for 16S rRNA gene amplicon sequencing. Variable region 4 (V4) of the 16S rRNA gene is currently the most commonly used marker for bacterial amplicon sequencing and the 515-806 fragment in particular has been shown to outperform alternative variable regions both in terms of reproducibility and classification accuracy [1,2]. The 515-806 fragment can be spanned by sequencing 200-300 nt from both ends, easily achieved on the Illumina MiSeq (which delivers 15-25 million paired reads). The use of a lengthy overlap between paired reads is necessary for reliable paired read merging, ensuring increased coverage on each sequence and higher quality data. Illumina’s HiSeq 2500 machine is capable of producing 300 million 250 bp paired reads per run using Rapid run mode v2 500 cycle reagents, at approximately one-third the cost per base compared to MiSeq.

Nucleotide diversity - having equal proportions of A, C, G, and T nucleotides at each base position in a sequencing library - is required for effective template generation on Illumina sequencing platforms [3]. In particular, the diversity in the first 11 bases of the DNA fragment is critical for cluster identification and color matrix estimation. Libraries with low complexity and uneven nucleotide distribution such as amplicon libraries can be successfully sequenced by blending in a balanced spike-in (Illumina recommends PhiX) which significantly reduces the amount of useful data obtained. This issue can be partially overcome by the addition of so-called heterogeneity spacers, short sequences of variable length, at the 5’ end of the amplicon thus introducing the required base diversity for effective sequencing with a minimal requirement for spike-in [4].

The novel protocol that we describe in this study adds a third Illumina compatible index to the existing dual indexing strategy developed initially by Kozich et al. [5]. We use a two-stage PCR protocol (as recommended by Illumina [6]), thus reducing the number of oligos required for multiplexing large numbers of samples. In addition, our protocol includes the heterogeneity spacers introduced by Fadrosh et al. [4] to increase nucleotide diversity at the start of sequencing reads, allowing greater utilization of available sequencing capacity.

Here, we benchmark our new sequencing protocol using both sequencing of a mock community and repeated sequencing of a standardized environmental sample. We looked for possible bias effects of PCR cycle number, input template DNA amount, PCR indexing of samples, MiSeq vs. HiSeq, between libraries and between runs variation. These types of evaluation are necessary to understand potential misinterpretations that could result from technical issues inherent in the design and practice of characterizing bacterial communities.

## Methods

Briefly, the procedure starts with a PCR reaction (PCR1) for general amplification of the V4 515-806 region of the 16S rRNA gene. Oligos used in this reaction includes the dual index sequence necessary for multiplexing (*see below*) along with heterogeneity spacers for increasing library complexity. Equal amounts of each amplified sample are then enriched in a normalization step and pooled. A second PCR reaction (PCR2) is then performed, using each amplicon pool as template, which primes on the ends of the oligos used for PCR1 (Fig. 1). This reaction adds a third index sequence and completes the Illumina adapters required for sequencing. Amplified PCR2 products are cleaned and quantified, then blended to make a single library before sequencing on the HiSeq 2500 (rapid mode) or MiSeq.

**Figure 1.**
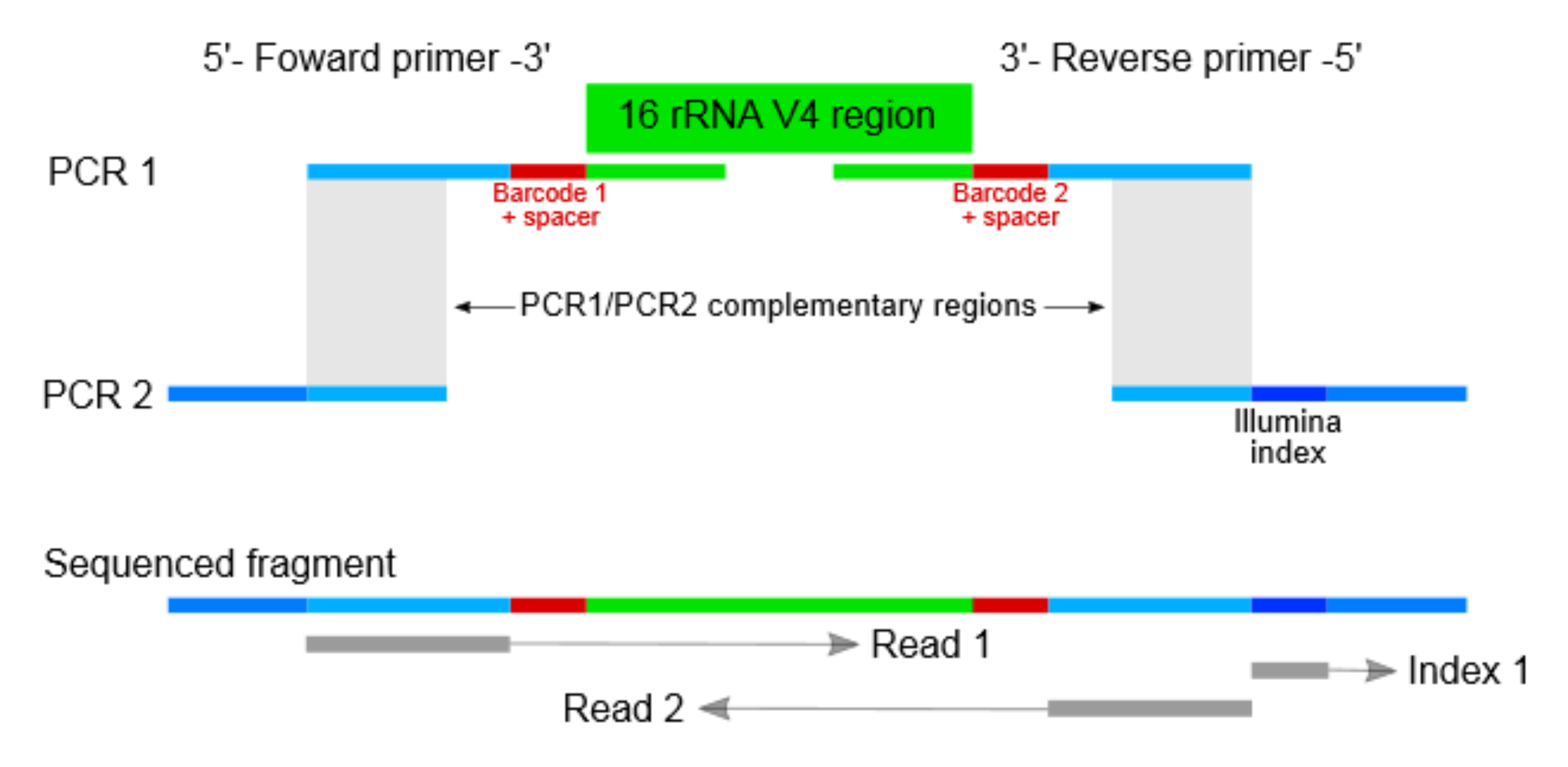
Triple indexing design. The triple indexing strategy incorporates two PCR steps. During the first PCR (PCR1) the template sequence of interest is targeted and amplified (green). The primers for this reaction also contain an indexing sequence and a heterogeneity spacer sequence (red), and a partial Illumina adapter (blue). A second PCR (PCR2) allows for the introduction of a third indexing sequence (dark blue) as well as completion of the Illumina adapter sequenc

### Oligo design

The primers used for PCR1 consists of four parts (Fig. 1): **1.** The 3’ end primes the V4 region of the 16S rRNA [7] with 515Fb and 806rB in forward and reverse primer, respectively. **2.** A heterogeneity spacer (0-7 bp) as described by Fadrosh, et.al [4]. **3.** Internal barcodes on both forward and reverse primers, also as described by Fadrosh et al. [4], to allow 96-384 sample index combinations. **4.** The 5’ end contains partial Illumina adaptors. The forward primer used in PCR2 is the same for all the samples and contains the remaining Illumina adaptor that is necessary for sequencing. Reverse primers contain the third index (6 nt Illumina TruSeq index) and the remaining Illumina adaptor. Complete lists of primers employed in this study are provided in (Additional file 1: Table S1: Not provided in this bioRxiv submission). All primers were purchased from Integrated DNA Technologies (Coralville, IA, USA) as 4nmol ultramers.

### Bacterial mock community

The mock bacterial community used in this study was generously provided by Prof. Lutgarde Raskin, Department of Civil and Environmental Engineering, University of Michigan and has been described elsewhere [8]. Briefly, near full length sequences of the 16S rRNA gene from 33 bacterial species representing more than 25 different phyla (Additional file 2: Table S2, Additional file 3: Figure S1) were cloned into plasmids. Plasmid concentrations were then measured before blending to equal proportions at a final concentration of 1x10e^9^ plasmids/μl.

### Library preparation

For an initial MiSeq sequencing run the PCR1 amplification was carried out with the following reaction mixture: 9μl H2O, 10μl 5Prime Hot Master Mix (Gaithersburg, MD, USA), 2.5μl of 1μM PCR1 forward primer, 2.5μl of 1μM PCR1 reverse primer, and 1 μl mock community template at either 2.5e^6^ or 2.5e^7^ molecules/μl. The following program was used for amplification: 94°C for 3 minutes for initial melting; 25, 30 or 35 cycles of 94°C for 10 sec, 50°C for 30 sec, 72°C for 45 sec; 72°C for 10 minutes. Products were verified on a 1% agarose gel, and all samples were cleaned and normalized to equal concentrations using the SequalPrep Normalization Plate Kit (Invitrogen, Carlsbad, CA, USA) according to the manufacturer's instructions. Samples assigned the same third index were pooled before the PCR2 reaction (Additional files 4-6: Tables S3-5). PCR2 amplification was carried out with the following reaction mix: 10μl H2O, 20μl 5Prime Hot Master Mix, 5μl of 1 μM PCR2 forward primer, 5μl of 1μM PCR2 reverse primer, 10μl template pool at 1 ng/μl. The following program was used for amplification: 94°C for 3 min; 5 or 10 cycles of 94°C for 10 sec, 58°C for 30 sec, 72°C for 45 sec; 72°C for 10 min. Products were again verified on a 1% agarose gel before cleaning with the Agencourt AMPure XP PCR purification kit (Beckman Coulter, Brea, CA, USA) according to the manufacturer’s recommendations. DNA concentration and fragment size were measured on a Qubit fluorometer (Invitrogen, Carlsbad, CA USA) and Agilent Bioanalyzer (Santa Clara, CA, USA), respectively. Samples with significant amounts of off-target fragment were cleaned one additional time with AMPure beads, but with the protocol modified to a 1:1 mixture ratio of beads and sample in order to eliminate shorter fragments. Libraries were blended in proportions to ensure similar coverage of all samples. For later HiSeq sequencing runs (Additional files 7-9: Tables S6-8) the library preparation procedure was identical to the MiSeq protocol with 35 and 10 cycles for PCR1 and PCR2, respectively, with the following exception: the repeated melting steps during both PCR amplifications were increased from 10 to 30 seconds.

### Illumina sequencing

Paired-end sequencing was performed on a MiSeq (Illumina, San Diego, CA, USA) using v3 reagents, generating 300 bp reads per end, or on a HiSeq 2500 (Illumina) in Rapid mode, with v2 reagents, producing 250 bp reads per end, according to manufacturer’s instructions. In each case, Illumina spike-in (PhiX) was included at 10% while loading was dropped to 80% normal recommended levels. Raw bcl files were processed using RTA v1.18.54 (MiSeq) and v1.18.66.3 (HiSeq). Initial de-multiplexing of the data based on the Illumina index reads (third index) was carried out using bcl2fastq v.2.17.1.14, which also converted the raw data to fastq files. Quality of the sequenced data was verified using FastQC v0.11.3 [9] and summarised using multiqc v0.3.1 [10].

### Downstream analyses of mock community data

Low quality reads were trimmed and Illumina adapters were removed using Trimmomatic v0.36 [11] with default settings. Reads mapping to the PhiX genome (NCBI id: NC_001422.1) using BBMap v36.02 [12] were also removed. De-multiplexing of data based on the dual index sequences was carried out using custom scripts. Internal barcodes and spacers were removed using cutadapt v1.4.1 [13] and paired reads were merged using FLASH v1.2.11 [14] with default settings. Sequence files were then converted from fastq to fasta and primers were trimmed from merged read ends. For each mock community sample, reads were aligned against a local database consisting of the 16S rRNA sequences of the 33 bacterial species included in the mock community using BLAST [15] and only reads with full length alignment with 100% sequence identity were retained for subsequent analyses.

The BLAST tabular output files were then imported into R [16]. OTU tables were produced by counting the number of occurrences of each bacterial species in the mock community. Bray-Curtis distance matrix computation, adonis modelling (Permutational Multivariate Analysis of Variance Using Distance Matrices) and ANOSIM (using Bray-Curtis dissimilarities and 10,000 permutations) were carried out using the R package vegan. Kruskal’s Non-Metric Multidimensional Scaling was carried out using the function isoMDS in the R package MASS with default settings. Chimera detection was carried out using the uparse [17] pipeline in usearch v8.1.1861 [18] with no minimum group size for the dereplication and clustering steps. Chimera formation rates were then found from the uparse output files.

### Environmental sample sequencing

A standardized environmental sample was made by suspending a fresh fecal sample (32.5g) from a healthy infant in 500ml of PBS. 1ml aliquots of this suspension were then transferred to microtubes and immediately frozen at -80°C. 500μl of sample material was used for DNA extraction with the PowerSoil 96 well DNA isolation kit (MO BIO Laboratories Inc., Carlsbad, CA, USA). A total of 54 standardized samples were DNA-extracted on 19 different plates and prepared as part of 19 different sequencing libraries (Additional file 8: Table S7), with 3 standardized samples per library with two exceptions (one library with two samples and one with only one). Library preparation and Illumina sequencing were carried out as described above, and the samples were sequenced as part of two separate HiSeq sequencing runs (26 samples in one, 28 in the other). Sequence processing was performed as described above up to and including paired read merging. Further processing was carried out using a combination of vsearch v2.0.3 [19] and usearch v8.1.1861 [18]. Specifically, dereplication was performed with the ‐‐ derep_fulllength function in vsearch with a minimum unique group size set to 2. OTU clustering, chimera removal, taxonomic assignment and OTU table building were carried out using the uparse pipeline [17] in usearch. Taxonomic assignment to the genus level was done against the RDP classifier. Read depths were normalized by common scaling [20]. This entails multiplying each OTU count in a given library with the ratio of the smallest library size (97,187 reads) to size of the library in question. This procedure replaces rarefying (i.e. random sub-sampling to the lowest number of reads) as it produces the library scaling one would achieve by averaging over an infinite number of repeated sub-samplings. After conversion of OTU counts to relative abundances, OTUs with an average relative abundance below 0.01% were removed from the data in order to reduce noise from artifacts.

We also sequenced 25 fecal samples from 5 healthy adults. We analyzed 4-6 samples, collected on consecutive days, per individual. With one exception the samples were prepared as part of the same library, and all samples were sequenced in one HiSeq run. Sequencing and subsequent analyses followed the exact same procedure as the standardized samples described above, with the exception that scaling was done against a smallest library size of 53,902 reads.

## Results

We evaluated several potential sources of bias in our sequencing strategy, using the total output from one MiSeq run and partial output from two HiSeq runs, resulting in six separate datasets (Table 1). Bias caused by sample indexing both in PCR1 (Dataset 1, Additional file 4: Table S3) and PCR2 (Dataset 2, Additional file 5: Table S4) reactions, as well as the effects of DNA template amount used for PCR and PCR cycle number (Dataset 3, Additional file 6: Table S5), were evaluated by sequencing the mock community 160 times in a single MiSeq run. We sequenced the mock community 24 additional times as part of a HiSeq run (Dataset 4, Additional file 7: Table S6) in order to assess design optimization, as well as to provide a basis for comparison between MiSeq and HiSeq results. Furthermore, a standardized environmental sample was sequenced 54 times as part of two separate HiSeq runs (Dataset 5, Additional file 8: Table S7) in order to assess variability between sequencing libraries and sequencing runs. Finally, we sequenced, in a single HiSeq run, 25 fecal samples collected on consecutive days from 5 adult volunteers (Dataset 6, Additional file 9: Table S8), in order to provide a baseline measure of variability in a realistic experimental setting, as well as to assess discriminatory power.

**Table 1.** Overview of the experimental design. Primer lists, read numbers and other pertinent information for Datasets 1-6 can be found in Additional files 4-9: Tables S3-8, respectively. Environmental samples were DNA extracted using the PowerSoil 96 well DNA isolation kit (MO BIO Laboratories Inc.) and processed as part of 19 separate sequencing libraries and two different HiSeq sequencing runs.

| <u>Dataset</u> | <u>Sequencer*</u> | <u>Evaluation</u> | <u>Data derived from</u> | <u>Primer list, read numbers and other information</u> |
| --- | --- | --- | --- | --- |
| 1 | MiSeq | PCR1 index | 96 mock community samples | Additional file 4: Table S3 |
| 2 | MiSeq | PCR2 index | 20 mock community samples | Additional file 5: Table S4 |
| 3 | MiSeq | PCR cycle number and input template amount | 48 mock community samples | Additional file 6: Table S5 |
| 4 | HiSeq | HiSeq protocol | 24 mock community samples | Additional file 7: Table S6 |
| 5 | HiSeq | Environmental samples 1 | 54 standardized fecal samples | Additional file 8: Table S7 |
| 6 | HiSeq | Environmental samples 2 | 25 fecal samples from 5 healthy adults | Additional file 9: Table S8 |
|  | *MiSeq - MiSeq v3600 cycle kit: 300 bp paired end run |  |  |  |
|  | *HiSeq - HiSeq 2500 rapid v2 500 cycle kit: 250 bp paired end run |  |  |  |

### Effect of PCR1 indexing

Twelve forward and eight reverse PCR1 primers were used in this design to generate 96 PCR1 index combinations (Dataset 1, Additional file 4: Table S3), i.e. a full standard plate. Since these primers contain different 12 bp indices, and have slightly different lengths due to the “heterogeneity spacer”, they can conceivably amplify with different efficiencies during PCR, which could bias results. To examine this effect, we amplified the same template (the mock community library, 2.5e^6^ input molecules per reaction) using 96 PCR1 primer combinations, employing 35 cycles of PCR1 and 10 cycles of PCR2 amplification (the same PCR2 forward and reverse primer was used for all reactions). These amplification conditions represent the maximum number of cycles (hence maximum possible bias) used in the study. The 96 samples were sequenced on the MiSeq platform yielding a mean of 80,330 (±31,559 s.d.) reads per sample after quality filtering and paired read merging.

Variation in yield from each PCR reaction can be expected due to amplification efficiency and technical variation caused by normalization prior to PCR2. Amplification efficiency was seen to be approximately equal from all reactions, as determined by running the products on an agarose gel (data not shown). However, index combinations where the same index sequence was present on both forward and reverse PCR1 primers clearly produced a lower yield of sequence data (Additional file 10: Figure S2), with mean read numbers at 8,968 (same index samples, n=8) vs. 86,818 (different index samples, n=88). However, apart from the relatively low read numbers, the species relative abundances measured in these samples did not differ noticeably from other samples, and thus were included in further analyses.

Based on the blending estimates described in Pinto and Raskin [8], the expected proportion of each species was 3%. Overall relative abundance estimates generally centered on the expectations (Fig. 2). Sequencing results from our experiment deviated from this expected value by 1.34% on average. However, some species deviated more than others from expectation. In particular, *Thermomicrobium roseum* was detected, at very low abundances, in only 7 of the 96 sequenced samples. Notably, this is the species with the highest GC content (69%) in the mock community, so low sequencing efficiency of the rDNA from this bacterium is unsurprising, as Illumina sequencing is not effective on sequences with extreme GC content [21]. Overall, significant spread in relative abundance estimates within species was observed, with a mean standard deviation of 0.66%. This could largely be attributed to bias effects from the primers used for PCR1 amplification causing significant structure in the data (Fig. 3), with 74.4% of the inter-sample variation in Bray-Curtis distances explained by samples having been amplified with different PCR1 reverse primers (p<0.001, ANOSIM) and 18.7% ascribed to the forward primers (p=0.003, ANOSIM), accounting for a total of 93.1% of observed variation.

**Figure 2.**
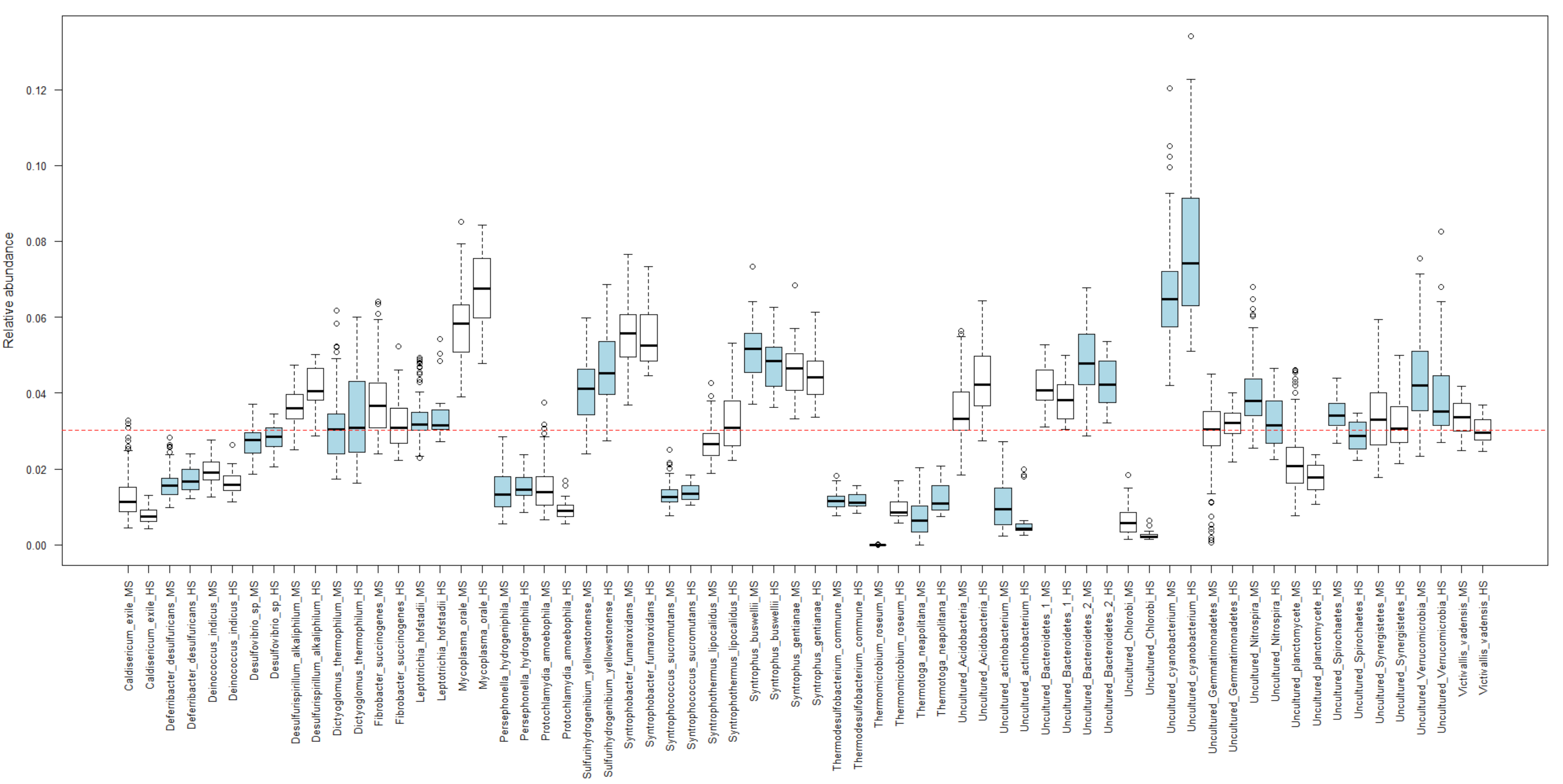
Relative abundances of the 33 bacterial species in the mock community sample estimated from both the MiSeq (Dataset 1, n=96) and HiSeq (Dataset 4, n=24) data (Additional file 4: Table S3, Additional file 7: Table S6). Species abundance estimates are shown side-by-side with MiSeq estimates labelled ‘MS’ and HiSeq estimates labeled ‘HS’. For enhanced visualization, each pair of colored bars (blue or white) depicts the estimated relative abundances for one species. The dotted red line shows the relative abundance expectation given perfectly equal blending. Each box represents the interquartile range while the whiskers represent 1.5 times the interquartile range. Points outside the whiskers represent outliers.

**Figure 3.**
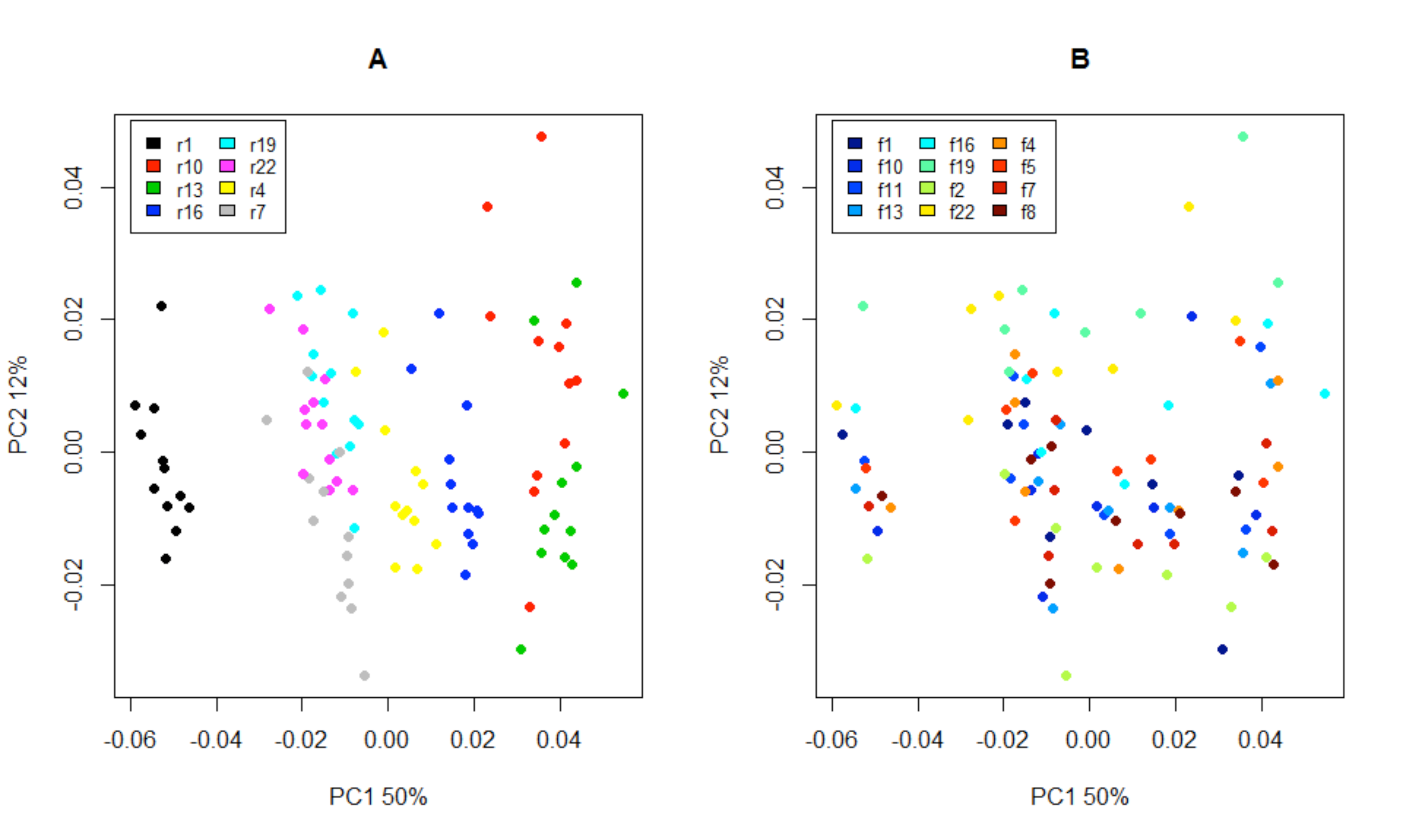
Scores plot based on a principal component analysis model computed from the matrix of species relative abundances from Dataset 1 (Additional file 4: Table S3). A. Samples are colored according to the reverse primer used for PCR1. B. Samples are colored according to the forward primer used for PCR1. In both A and B, the first two dimensions, explaining 62% of the total variance, are shown.

Observed effects were primarily associated with the oligo indices designated r1, r10 and r13 (Fig. 3A). These effects were again associated with specific bacterial species (Additional file 11:Figure S3), e.g. r1 appeared to be biased in favor of *Mycobacterium orale* and an uncultured *bacteriodetes* while r10 and r13 favored an uncultured *cyanobacterium* and an uncultured *verrucomicrobium*. In general there was good agreement between the 96 measured mock communities with an average Spearman correlation of 0.85 between the vectors describing relative species abundances, but the relatively poorer agreement between samples processed with the r1 primer and those amplified with the r10 and r13 primers was evident, with correlations even falling below 0.5 (Additional file 12: Figure S4). We also observed a significant negative relationship between estimated mean relative abundances and GC content of sequenced fragments (p=0.002, linear regression; Fig. 4), with relative abundance decreasing by 0.18% for each one percent increase in GC content. Chimera formation rates, at a mean of 17% (±2.2 s.d.), were not significantly associated with specific oligo combinations, except for a slightly elevated rate for samples amplified with primer r13 (p<0.001, linear model, (Additional file 13: Figure S5). Furthermore, it was evident that the large majority of chimeric sequences occurred only once in any given sample, with a mean proportion of singleton chimeras at 90%.

**Figure 4.**
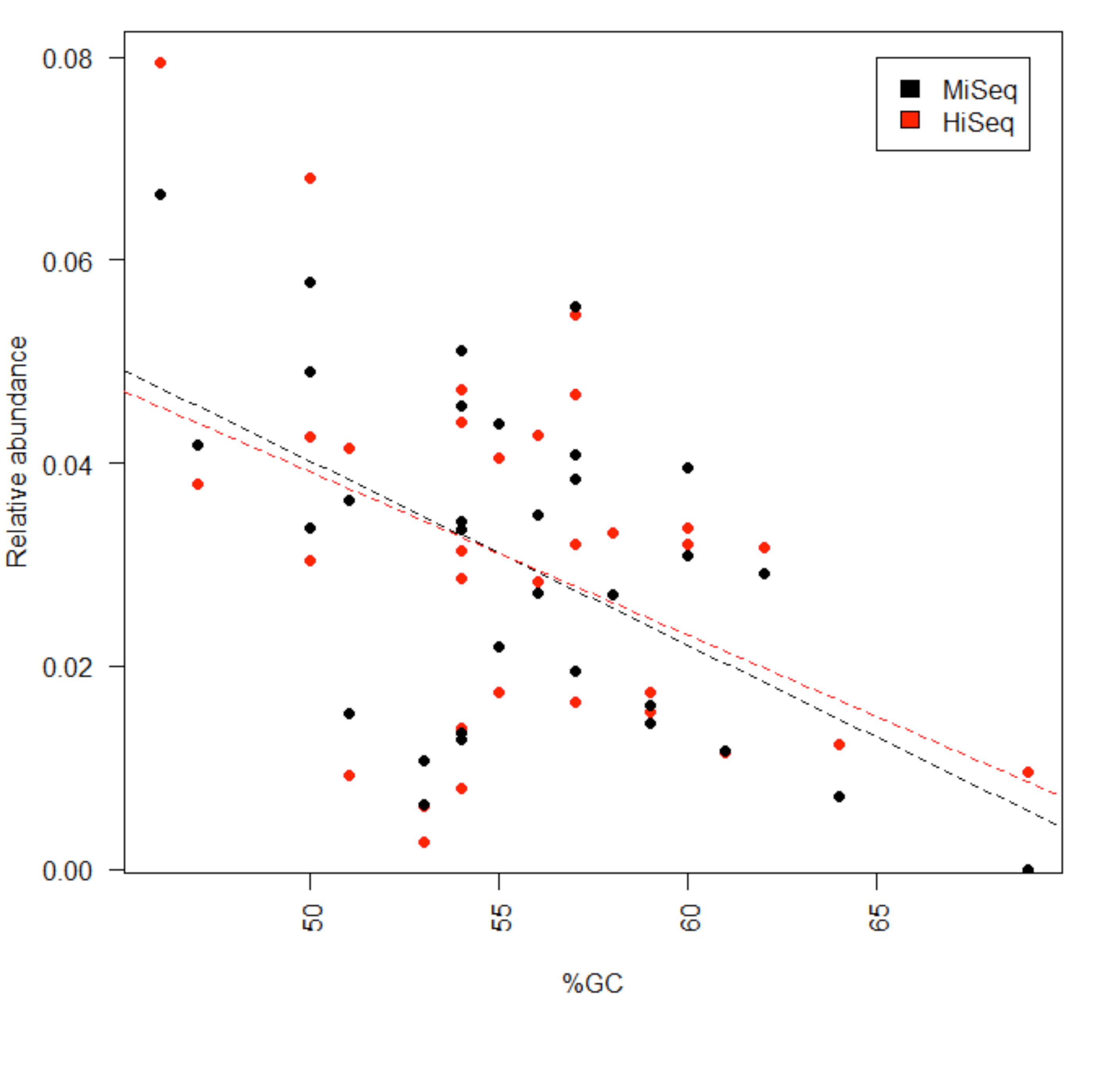
Relationship between mean relative abundance estimates and GC percentage for Datasets 1 and 4. There is a significant negative linear relationship for both the MiSeq (p=0.002, n=96, Additional file 4: Table S3) and HiSeq (p=0.012, n=24, Additional file 7: Table S6) data. Estimates drop by 0.18% and 0.16% for each 1% increase in GC content for the MiSeq and HiSeq estimates, respectively.

### Effects of PCR2 indexing

Similar to the analysis above, we attempted to quantify bias emerging from the use of different index sequences introduced in PCR2 amplification. In the case of PCR2, however, indices were only present on the reverse primer, were shorter (6 bp), and no heterogeneity spacers were required. Furthermore, the priming sequence used for this amplification did not contain wobble-bases (Additional file 1: Table S1: Not provided in this bioRxiv submission), further reducing the potential for bias. We compared relative species abundances measured in five sets of PCR1 products (four per set for a total of 20 samples) amplified using 35 cycles, with each set of four samples using a different PCR2 index primer (Dataset 2, Additional file 5: Table S4) and 10 cycles of amplification. The samples were sequenced on the MiSeq platform yielding and average of 144,171 (±100,253 s.d.) reads. PCR2 was again performed for 10 cycles of amplification, to introduce the maximum possible bias. By far the largest proportion of intersample variation in Bray-Curtis distances was explained by differences caused by PCR1 reverse priming oligos (82.7%, Adonis model), with the main difference between samples amplified using r1 and r10 (Additional file 14: Figure S6). Conversely, only 8.6% (Adonis model) of the observed distance variation could be related to differences between oligos used for PCR2 amplifications, and these effects were not found to be statistically significant (p=0.98, ANOSIM).

### Effects of PCR cycle number and template amount

PCR has on numerous occasions been shown to introduce amplification bias [22]. We therefore wished to examine how the number of amplification cycles, during PCR1 and PCR2, affected sequencing results. To this end we sequenced 48 mock community samples (MiSeq, mean of 104,705 (±43,026 s.d.) reads per sample) according a factorial design (Dataset 3, Additional file 6: Table S5). Briefly, we performed PCR1 amplification using either 25, 30 or 35 cycles (16 samples per treatment). To investigate the effect of PCR2 cycle number, PCR1 products were further amplified with either 5 or 10 cycles (24 samples per treatment). Furthermore, within each combination of PCR cycle number regimes, initial PCR1 amplification was carried out using either relatively low (2.5e^6^ molecules) or high (2.5e^7^ molecules) amounts of DNA template (24 samples per treatment). This design gave a total of four samples within each possible combination of variable levels. We did not observe any significant interaction effects between template amount and PCR cycle number (p=0.94 and 0.98 for PCR1 and PCR2, respectively, Adonis models), or between PCR1 and PCR2 cycle regime (p=0.82, Adonis model). Thus, for further statistical analysis each variable was treated as separate from the two others, with the number of samples per treatment as stated above.

Significant effects of input DNA template amount for the PCR1 amplification were observed (p<0.001, ANOSIM; Fig. 5), particularly related to the species *Fibrobacter succinogenes* being favored in reactions using the relatively high template concentration (Additional file 15: Figure S7). However, the abundance estimates were remarkably robust to the effect of DNA input amount over the 10-fold range tested here, explaining only 12.1% of the total observed variation in inter-sample Bray-Curtis distances (Adonis model).

**Figure 5.**
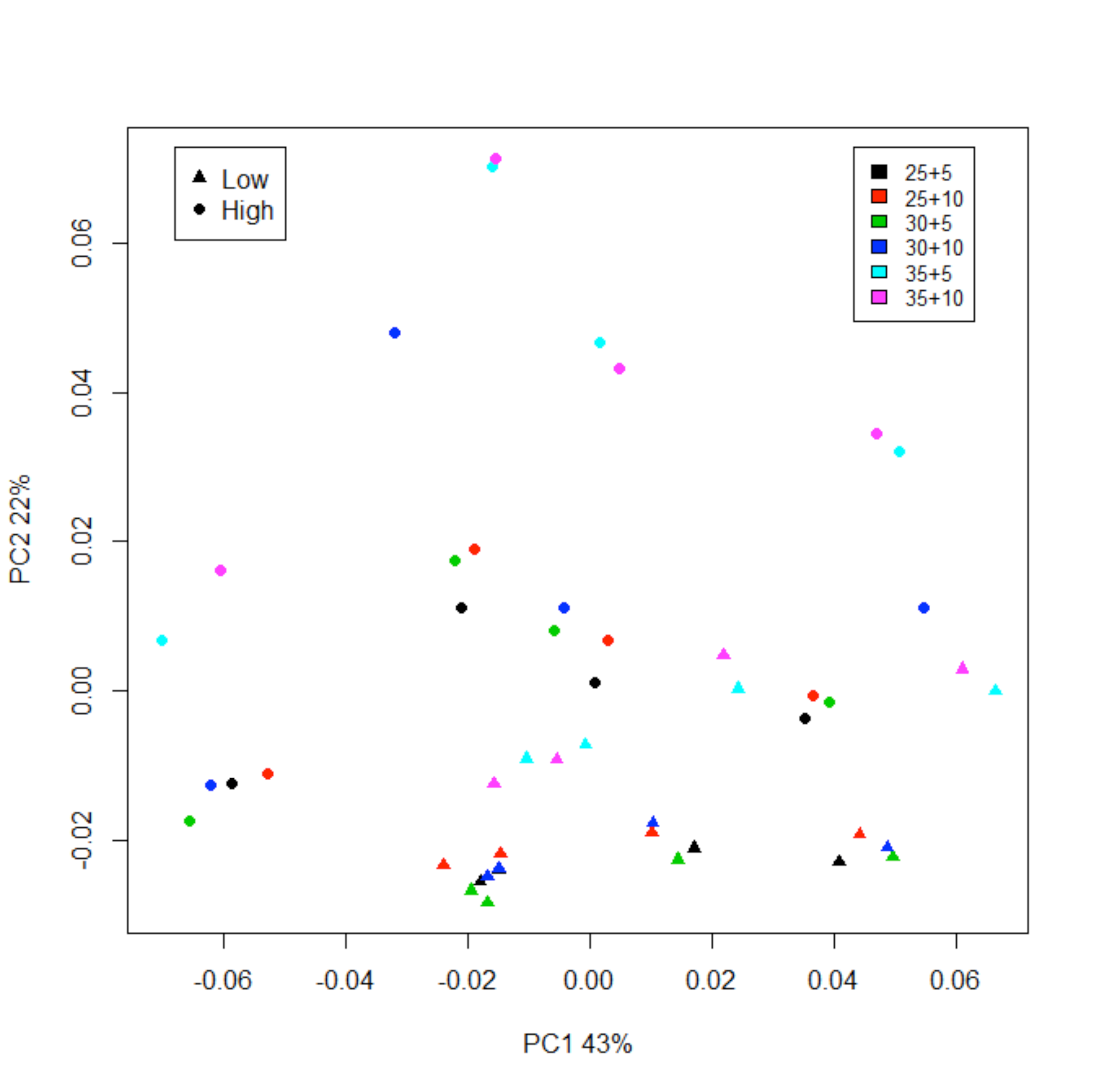
Scores plot based on a principal component analysis model computed from the matrix of species relative abundances from Dataset 3 (Additional file 6: Table S5). Samples are colored according to PCR1 and PCR2 cycle regime, with the number of cycles indicated in the legend (PCR1 + PCR2). Filled dots and triangles represent samples prepared with tenfold difference in input DNA template concentration used for PCR1. The first two dimensions, explaining 65% of the total variance, are shown.

There was a significant, albeit noisy, effect of the number of PCR1 cycles on the number of sequence reads produced (Additional file 6: Table S5), with read numbers on average increasing with 4084 reads per added cycle (p=0.006, linear regression). No corresponding effects were observed for PCR2 cycle number or template amount (p=0.91 and 0.89, respectively; linear models). We observed a clear but subtle effect of the number of cycles used in PCR1 reactions on estimated relative abundances (p<0.001, ANOSIM) (Additional file 16: Figure S8), with this factor accounting for 13.6% of the observed variation in Bray-Curtis distances (Adonis model). However, comparisons of squared deviations from the expected relative abundances between samples that had undergone different numbers of PCR1 cycles did not find any particular regime to produce results significantly closer to the expectation (25 vs. 30 cycles: p=0.45; 25 vs. 35 cycles: p=0.30; 30 vs. 35 cycles: p=0.86, unpaired t-tests). All in all there were 10 species for which we observed a significant linear relationship between measured relative abundance and the number of cycles used during PCR1 amplification (Fig. 6). PCR1 cycle number seemed to particularly affect abundance estimations of species with a high GC content, e.g. *Thermotoga neapolitana* and *Thermodesulfobacterium commune* (GC of 64 and 61%, respectively). One extreme example is *Thermomicrobium roseum*, the species with the highest GC content (69%) on the sequenced fragment. This species was observed in 15 out of 16 sequenced samples, albeit at very low abundances, when using 25 PCR1 cycles, but only in 5 out of 32 samples at higher cycle numbers (relative abundances <0.01%). Another noteworthy example is the uncultivated gemmatimonadetes species (GC of 62%) which was readily observed after 25 and 30 PCR cycles, but dropped sharply at 35. Conversely, we observed a positive relationship for four species of low or intermediate GC content (Figure 3). The number of cycles used for PCR2 did not produce discernible effects (p=0.84, ANOSIM) (Additional file 17: Figure S9) and accounted for only 1.1% of observed distance variation between samples.

**Figure 6.**
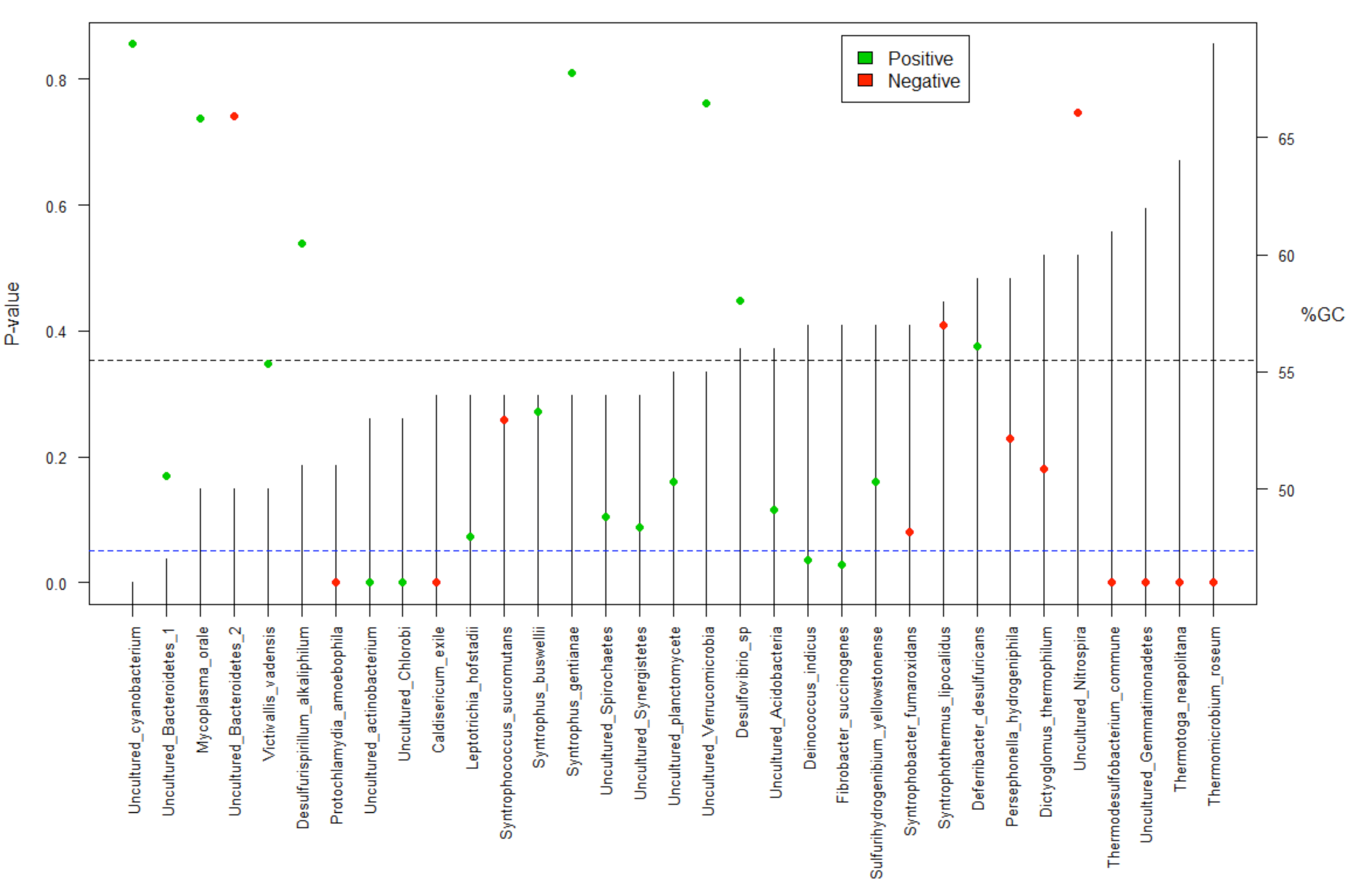
Statistical significance and direction of relationships between estimated relative abundances of sequence reads and PCR cycle number (Additional file 16: Figure S8) in Dataset 3. The dots represent p-values from linear regression models, with green and red representing positive and negative relationships, respectively. The species are ordered according to the GC content on the sequenced fragment (vertical lines). The dotted blue lines signifies the significance threshold of p=0.05 (left axis), while the dotted black line represents the mean GC percentage (right axis).

Even when considering the bias sources built into the experimental design in terms of PCR regime and template amount, the by far strongest effects on data structure could be ascribed to bias related to the primers used for PCR1 amplification. As noted above, the DNA oligos used for forward and reverse priming differed only in their respective index/spacer sequences, yet 62.1% of inter-sample variation in Bray-Curtis distances was explained by samples having been amplified with different reverse primers during PCR1. As above, the forward primers had a much smaller effect, contributing only 2.1% of inter-sample variation. All in all, 91.1% of the total distance variation could be explained by the combination of PCR cycle number effects, template concentration and PCR1 priming bias.

### Chimeric read formation

Chimeric amplification artifacts, arising from two or more templates, artificially inflate diversity estimates in 16S sequencing data. Over-amplified libraries can include up to 45% such artifacts [23]. We detected a strong linear association between PCR cycle regime and chimeric sequence formation when considering PCR1 cycle numbers, with the proportion of chimeric sequences increasing with as much as 1% for each added reaction cycle (R^2^=0.81, p<<0.001, linear model)(Additional file 18: Figure S10). We did not observe a significant relationship when only considering the second step PCR2 regime (p=0.61, t-test), however when considering the combined effects of both PCR1 and PCR2, there was still a strong linear relationship (R2=0.65, p<<0.001, linear model: Fig. 7). It was also evident that for samples having undergone an equal total number of PCR cycles (PCR1 +PCR2) partitioned differently between first and second step reactions, i.e. 25+10/30+5, and 30+10/35+5, the mean chimera formation rates were significantly higher in the samples having undergone more first step cycles (p<0.001 for both comparisons, t-tests). We did not observe significant effects of input template amount on chimera formation rates (p=0.72, t-test).

**Figure 7.**
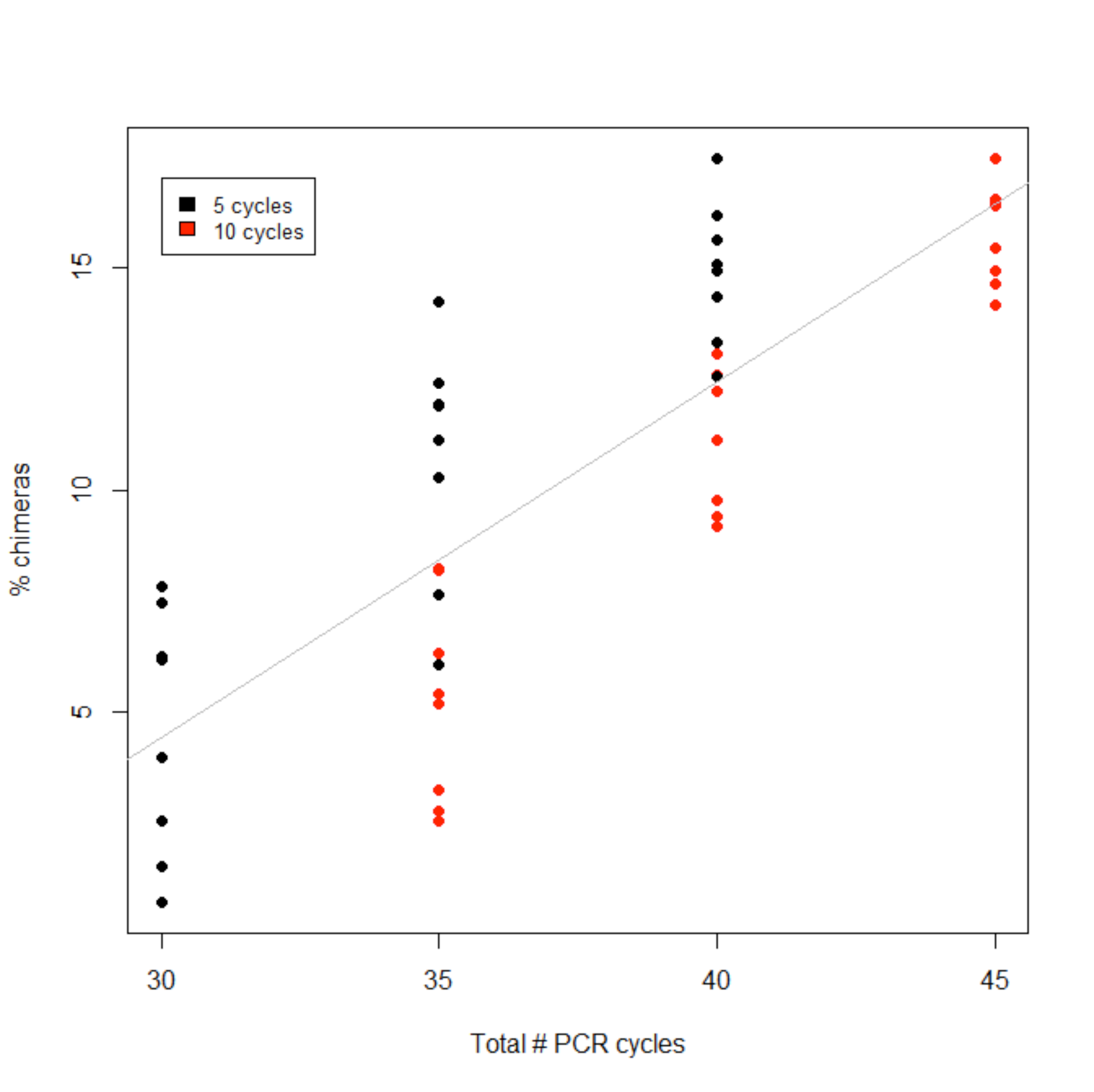
Relationship between PCR cycle number and chimeric sequence formation in Dataset 3. The combined numbers of PCR1 and PCR2 amplification cycles are indicated on the x-axis. Black and red dots indicate samples amplified using 5 and 10 cycles for PCR2 respectively. A highly significant linear relationship (p<<0.001, linear regression model) was observed. The effects were primarily related to the PCR1 cycle number, e.g. samples undergoing 35 cycles (25 cycle PCR1 and 10 cycles PCR 2) had less chimeras than samples undergoing 35 cycles (30 cycles PCR1 and 5 cycles PCR2).

### Optimization and scalability to the HiSeq platform

In order to test the scalability of our protocol, and to investigate the potential for increasing measurement consistency by reducing bias introduced through sub-optimal primer combinations, we sequenced the mock community 24 more times (Dataset 4, Additional file 7: Table S6) as part of a HiSeq rapid run producing 250 bp paired-end reads (mean read number of 297,930 (±91,387 s.d.)). In this experiment we eliminated any instance of identical indices on the forward and reverse primers, and we replaced the poorly performing primers r1, r10 and r13 with alternative ones (r3, r12 and r15; Supplementary table S6). These primers were chosen because they have similar heterogeneity spacer lengths to the ones they replaced (Additional file 1: Table S1: Not provided in this bioRxiv submission). We also extended the melting time during PCR from 10 to 30 seconds to try to alleviate bias caused by high the GC content of some fragments.

Overall measurement accuracy was comparable with the MiSeq run described in the previous experiment (Fig. 2) with a mean deviation from the expected abundances at 1.38%, and observed differences in estimated community composition were not statistically significant (p=0.84, ANOSIM). Significant structure caused by bias both on the reverse and forward primers was still observed (p<0.001 and p=0.01, respectively, ANOSIM), with 71.9% and 23.1% (Adonis model) of the observed inter-sample variation in Bray-Curtis distances explained by the reverse and forward priming oligos used for PCR1 amplification, respectively (Additional files 19 and 20: Figures S11 and S12). The most severe discrepancies were caused by the new primer combinations (r3, r12 and r15), suggesting that spacer length may be playing a part in amplification bias. However, the new primers performed better than the indices that were replaced, as Spearman correlations in this run did not go below 0.66 (Additional file 21: Figure S13). The mean spearman correlation of the estimated OTU composition vectors was 0.87. The bacterial species that were subject to the most severe bias were the same as before (Fig. 2). Importantly, in this sequencing run we were consistently able to observe, at moderate abundances, the *Thermomicrobium roseum* sequences that were absent from the MiSeq run under the same 35 cycle PCR1 and 10 cycle PCR2 regimes with a mean relative abundance increase from 0.00023% to 1.0%. Estimates for other high GC content species *Thermotoga neapolitana* and uncultured *gemmatimonadetes* species also saw moderate increases in mean relative abundance (0.7%-1.2% and 2.9%-3.2% respectively). We attribute this to the increased time for DNA denaturation used for both PCR1 and PCR 2 amplification prior to sequencing on the HiSeq, and this result suggests that denaturation time during PCR can affect measurements, especially for high GC content species.

### Sequencing of environmental samples

Further evaluation of the sequencing protocol was done by sequencing human fecal samples. First, a standardized sample was sequenced 54 times using a variety of indices (Dataset 5, Additional file 8: Table S7). These samples were processed as part of nineteen different rounds of DNA extraction and subsequent library preparation (i.e. on 19 different 96 well plates; Table 1). Furthermore, they were sequenced as part of two different HiSeq runs, with 26 samples sequenced on the first run and 28 on the second. The mean Bray-Curtis distance of estimated bacterial community composition between samples was 0.18, a value slightly higher than that observed for the mock community (0.12) (Fig. 8). Most of the variation (54.4%, Adonis model) in inter-sample distances could be ascribed to batch effects (p<0.001, ANOSIM), i.e. the fact that the samples were processed on different plates. This could be due to differences in DNA extraction efficiency or downstream steps of library preparation. 20.1% of the variation could be attributed to amplification bias related to the reverse primers used during the PCR1 amplification (p<0.001, ANOSIM), while 3.7% of the variation was related to the forward primers (p=0.009, ANOSIM). 14% of the variation could be explained by the samples having been sequenced on two separate HiSeq runs (p<0.001, ANOSIM). In total, these four factors accounted for 92.3% of the observed variation.

For the purpose of comparison, 25 fecal samples were obtained from five healthy adult volunteers. The samples were taken sequentially on a daily basis (4-6 days), and sequenced on a single HiSeq run (Dataset 6, Additional file 9: Table S8). The purpose of this was to provide a baseline measure of inter-sample differences, as well as to evaluate relative effects of bias caused by indexed PCR, in a realistic experimental setting. The mean Bray-Curtis distance of this sample set was 0.57, illustrating that the variation observed in the data from both the mock community and the standardized fecal sample are comparatively low (Fig. 8). The inter-sample variation in distances could in this instance mainly be ascribed to differences between the individuals from which the samples originated (74.1%, Adonis model), while only 11.0% and 8.7% of the total variation was explained by the reverse and forward PCR1 primers, respectively (93.8% explained by all three factors). This demonstrates good potential for detecting biological signal in the face of inherent technical bias (Additional file 22: Figure S14).

**Figure 8.**
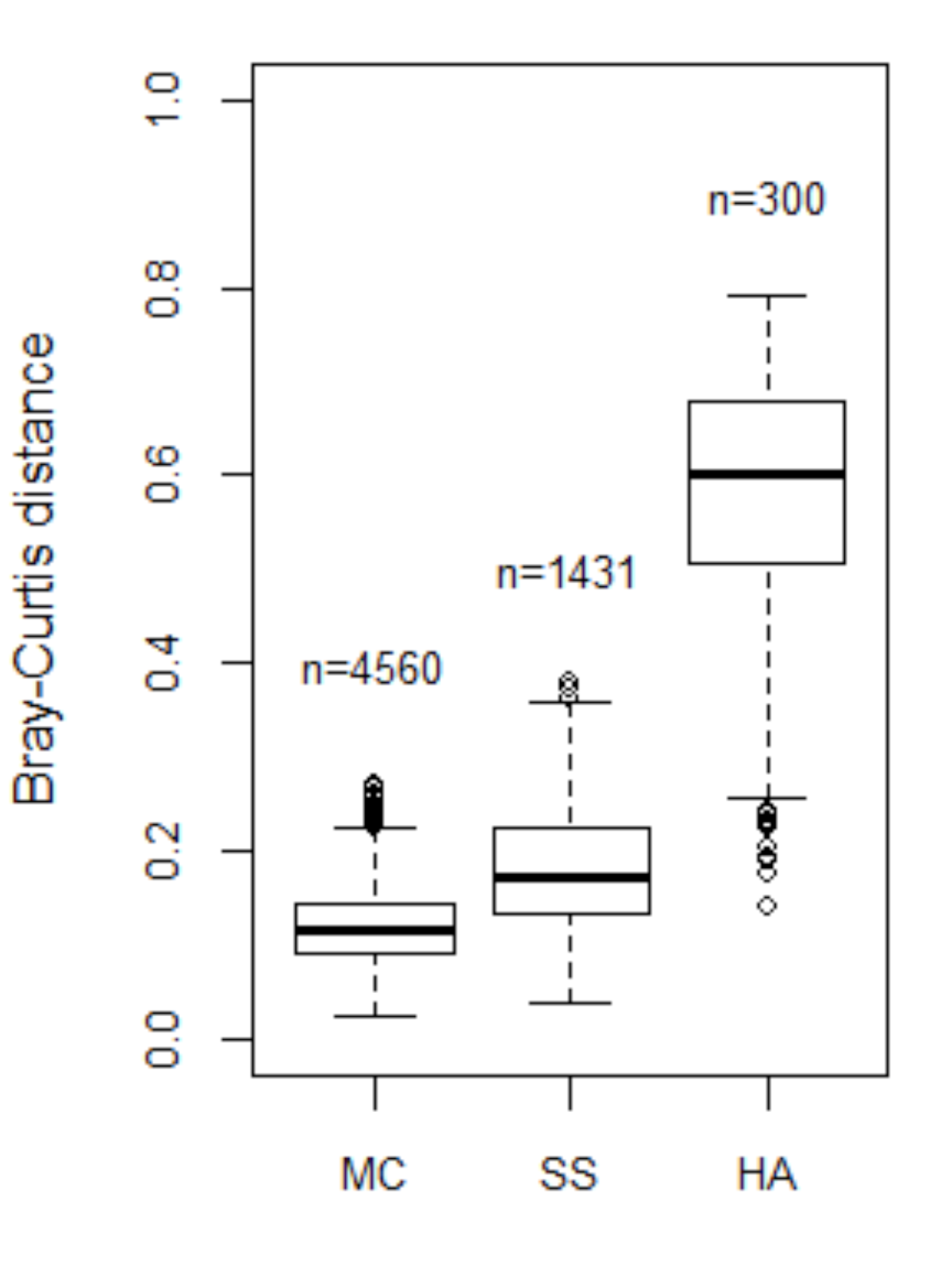
Pairwise Bray-Curtis distances for the mock community (MC, Dataset 1, Additional file 4: Table S3), standardized sample (SS, Dataset 5, Additional file 8: Table S7), and healthy adult (HA, Dataset 6, Additional file 9: Table S8) group. Each box represents the interquartile range while the whiskers represent 1.5 times the interquartile range. Points outside the whiskers represent outliers. The number of pairwise distances for each group is indicated over the boxes.

### Discussion

In this study we carried out extensive benchmarking of an Illumina sequencing library preparation protocol incorporating several extant features in order to improve on popular designs. Our main recommendations for optimal use of the protocol described here are summarized in Box1. The use of a triple indexing strategy allows for extensive multiplexing with a greatly reduced requirement for long custom oligos. This approach is also in theory extendable to quadruple indexing, with the addition of a barcode sequence on the reverse primer used during PCR2. Addition of heterogeneity spacers [4] allowed for PhiX spike-in at low levels while maintaining high data quality. Finally, the rapid run mode available on the Illumina HiSeq 2500 allows for a read length that easily spans the ~300 bp of the V4 marker fragment when using paired-end sequencing, and generates up to ten times as much data as a MiSeq run, allowing faster data generation on a single machine run.

The original article analyzing the mock community used in this study [8] reported significant deviations from the expected relative abundances according to their blending design. It should be noted that plasmid blending represents a significant error source, and some deviation from the intended ratios is inevitable. Pinto and Raskin [8] reported poor detection of some species, such as the thermophiles *T.neapolitana* and *T.commune*. These species were also detected below expected abundances in our study. On the other hand the previous study reported very low recovery of sequences from *V.vadensis* and *M.orale*. We detected *V.vadensis* at around the expected 3%, while *M.orale* was highly overrepresented in our data at around 6%. Conversely, Pinto and Raskin [8] found sequences from the two *Syntrophus* spp. to be overrepresented in their data, as did we. However, they detected the uncultured *planctomycetes* far over the expected 3% while we found it to represent around an average of 2% of reads. Direct comparison between the two studies is complicated by the fact that they used primers targeting different regions of the 16S rRNA gene, a known cause of differential PCR amplification bias [2]. Furthermore, we employed the Illumina sequencing platform as opposed to 454 pyrosequencing, which was used in the older study.

The input template DNA amount used for PCR1 had only moderate effects on the estimated relative abundances in the mock community, and the effects we did observe were consistently associated with specific species. In particular, for *Fibrobacter succigenes*, a major cellulolytic component of the ruminant microbiota [24], we observed an increase from 3.3% to 6.7% mean relative abundance when going from low to high amounts of template DNA. It is unclear whether this type of bias is specific to the mock community we analyzed, or if it represents a more general phenomenon. Wu et al. [25] sequenced various dilutions of sediment samples without observing significant effects on estimated community composition. This suggests that template amount bias might be system specific and should be treated on an *ad hoc* basis. By sequencing dilutions of a subset of samples within a given experiment, one could determine if bias is associated with particular taxa, and either error correction could be employed or caution used when making conclusions based on taxa that are more prone to template amount bias.

It is well established that multi-template PCR is subject to several sources of bias [26, 27], and these error sources can be aggravated when using indexed primers for highly multiplexed DNA sequencing [28]. The PCR cycle number used for amplification of a target fragment is a factor that can be expected to have effects on amplicon sequencing results. As expected, the number of PCR cycles used for amplification had a big impact on chimeric sequence formation rates, in line with previous findings [2]. Interestingly, chimera formation was much more strongly associated with the PCR1 reactions than PCR2 reactions (Fig. 7), indicating that primer degeneracy or the appended indices and adapters, or a combination of the two, increases the probability of template switching during PCR amplification relative to an amplification with perfectly matching primer sequences, such as for the PCR2 reactions used in our protocol. The high proportion of singleton chimeric reads (90%) also suggests that simple abundance filtering can greatly help towards elimination of chimeric sequences from the dataset. The remainder can be eliminated using one of several available chimera detection programs [17, 23].

We only observed relatively subtle effects of PCR1 cycle number and estimated relative abundances of species in the mock community. There was, however, a systematic negative relationship between PCR cycle number and estimated relative abundances of sequences with very high GC content (Fig. 6). This is a phenomenon that has been observed previously [29]. A likely explanation is that the relatively higher melting temperatures of these sequences causes them to be more easily outcompeted by lower GC containing template sequences due to differential amplification efficiency. Increasing the denaturation time during PCR1 increased recovery of the high GC content sequences in our mock community (*Thermomicrobium roseum, Thermotoga neapolitana* and uncultured *gemmatimonadetes*). The effects of melting time during PCR should be especially considered for bacterial communities expected to containing species with a wide range of GC contents. We also observed a significant negative relationship between estimated mean relative abundances and GC content of sequenced fragments in general. This suggests that a GC correction factor could be employed where relative abundance estimates are multiplied by some empirical constant in order to compensate for high GC content. The exact value of the correction factor (in our case 0.18% per percent increase in GC) should be corroborated by further experimentation.

Unexpectedly, we observed greatly reduced sequencing output for samples using identical index sequences on the forward and reverse primers during PCR1 (a reduced output by a factor of around 10 was observed in all 15 cases). A cursory analysis suggests that this phenomenon is caused by formation of high melting temperature hairpin structures that interfere with sequencing efficiency.

Primer indexing was found to be an important structuring factor affecting relative abundance estimates. This has been observed by other investigators [8, 28, 30]. The bulk of the observed variation (74.4%) due to primer indexing in this study was caused by the reverse primers. We were able to partially alleviate this problem by replacing underperforming primers (Additional file 1: Table S1: Not provided in this bioRxiv submission), with the total amount of the variation ascribed to primer indexing decreasing from 93.1% to 79.7%. Some bias caused by primer indexing during multi-template PCR is inevitable; however, an informed rationale for designing primer indices/spacers that minimize bias would represent a significant advancement. Possible ways of reducing this source of bias are by first using an unindexed PCR followed by either ligation of index and adapter sequences [2] or by using a second step indexed PCR [28]. We did not observe significant bias due to the non-degenerate PCR2 amplifications (which used six-nucleotide indices regularly employed by Illumina library preparation reagents on the reverse primers), suggesting that carrying out indexing PCR on amplicons, rather than genomic DNA, may alleviate primer indexing bias.

Primer indexing, although significant, was a relatively minor contributor to the overall variation when sequencing a standardized human fecal sample. In this case, the main variation (54.4%) in inter-sample distances was related to batch effects, such as DNA extraction being performed on different days. This highlights the importance of having highly standardized protocols starting with DNA extraction and through to the actual sequencing step. Although the mean Bray-Curtis distances between samples were higher for the standardized fecal samples than the mock community (Fig. 8), the mean spearman correlation of the estimated OTU composition vectors of both the mock community (0.85) and the standardized fecal sample (0.87) were similar. This suggests that one should get consistent results if one chooses either to benchmark by sequencing a defined mixture of 16S rRNA molecules or by repeated sequencing of a standardized environmental sample. Comparison with fecal samples taken sequentially from five adults demonstrated that the inter-sample distances observed in a real-world experimental setting are significantly higher than observed in both the mock community and standardized environmental sample data. In this experiment, primer indexing bias was only a minor contributor to the overall variation, while the biological signal resulting from sample origin accounted for 74.1% of the total observed variation. Our results highlights that even faced with a myriad of error sources, amplicon sequencing has a tremendous potential for providing detailed quantitative descriptions of complex microbial communities.

##### Box 1:Recommendations

- Highly standardized DNA extraction and sample processing protocol
- 30 second denaturation step for both PCRs
- Use different index on forward and reverse primers in PCR1
- Normalize dual indexed samples
- Library pooling after first PCR
- 10 cycles in PCR2
- Final cleanup (1:1 AMPure for size selection)
- Final library pooling of triple indexed samples
- 250 bp paired end sequencing in HiSeq 2500 Rapid run mode

## List of abbreviations

rRNA: Ribosomal RNA
PCR: Polymerase chain reaction
OTU: Operational taxonomic unit
BLAST: Basic local alignment search tool
MDS: Multidimensional scaling
PBS: Phosphate buffered saline
ANOSIM: Analysis of similarity
GC content: Guanine-cytosine content

## Declarations

### Ethics approval and consent to participate

All work with samples of human origin in this study has been approved by the Regional Ethics Committee of South-East Norway (REK nr. 2014/656). Consent for participation was obtained from all individuals involved in the study.

### Consent for publication

Consent for publication was obtained from all individuals involved in the study.

### Availability of data and materials

Amplicon sequencing datasets will be uploaded to European Nucleotide Archive prior to the publication of the manuscript. Authors will start the process of uploading the data shortly.

### Authors’ contributions

EJD, PT, GDG, and AYMS designed and carried out the experiments and wrote the paper. EJD and PT performed the statistical analyses.

## Acknowledgements

The authors would like to acknowledge Anders Ass for useful discussions. We would also like to acknowledge the voluntary participants for providing sample material.

## Additional file legends

Additional File 1: Supplementary Table 1: Not provided in this bioRxiv submission
**Filename: Additional_File_1-Table_S1_oligos.xls**
**Title: List of PCR primers used in this study.**
The designation ‘f’ or ‘r’ is used throughout the study to distinguish between forward or reverse primers respectively. The complete sequence of each oligo is given, and also the sequence of the individual elements of each oligonucleotide separately (partial Illumina adapter, index sequence, spacer sequence, and priming regions of v4 16S rRNA gene).

Additional File 2: Supplementary Table 2
**Filename: Additional_File_2-Table_S2_GC_content_33species.xls**
**Title: Taxonomy of bacteria in the mock community.**
List of bacterial species for which full length bacterial 16S rRNA gene sequences were cloned and included in approximately equal proportions in the mock community. Species are listed in the first column and GC content is listed in the next.

Additional File 3: Supplementary Figure 1
**Filename: Additional_File_3-Figure_S1_phylogeny.pdf**
**Title: Phylogeny of bacteria in the mock community.**
Neighbor joining tree showing the phylogenetic relationship among the 33 species represented in the mock community used for method validation. The tree is based on the 515-806 fragment (V4 region) of the 16S rRNA gene. The scale bar at the bottom refers to the number of pairwise nucleotide differences.

Additional File 4: Supplementary Table 3
**Filename: Additional_File_4-Table_S3_library1.xls**
**Title: Primers and read numbers for Dataset 1.**

Additional File 5: Supplementary Table 4
**Filename: Additional_File_5-Table_S4_library3to7.xls**
**Title: Primers and read numbers for Dataset 2.**

Additional File 6: Supplementary Table 5
**Filename: Additional_File_6-Table_S5_library2.xls**
**Title: Primers, read numbers, template amount and PCR cycle numbers for Dataset3.**

Additional File 7: Supplementary Table 6
**Filename: Additional_File_7-Table_S6_HS_mock.xls**
**Title: Primers and read numbers for Dataset 4.**

Additional File 8: Supplementary Table 7
**Filename: Additional_File_8-Table_S7_SS_samples.xls**
**Title: Primers, read numbers, run number and library numbers for Dataset 5.**

Additional File 9: Supplementary Table 8
**Filename: Additional_File_9-Table_S8_HA.xls**
**Title: Primers, read numbers and individual numbers for Dataset 6.**

Additional File 10: Supplementary Figure 2
**Filename: Additional_File_10-Figure_S2_readcounts.bmp**
**Title: Read counts for all PCR1 primer combinations in Dataset 1.**
Each bar represents the read count of a mock community sample after quality filtering and paired read merging. The figure shows data for all 96 PCR1 index combinations (Dataset 1, Additional file 4: Table S3). Tenfold fewer reads were observed in samples that were amplified with primer pairs using the same forward and reverse indices.

Additional File 11: Supplementary Figure 3
**Filename: Additional_File_11-Figure_S3_loadings_96sample.bmp**
**Title: Principal components analysis loadings plot showing the bacterial species defining the two largest variance components in Dataset 1.**
The main axis of variation accounts for 50% of the total and defines the difference between samples amplified with PCR1 primers r1 (e.g. *Mycoplasma orale*) and r10 and r13 (e.g. uncultured *verrucomicrobium*).

Additional File 12: Supplementary Figure 4
**Filename: Additional_File_12-Figure_S4_correlation_map_L1.bmp**
**Title: Heat map of pairwise sample correlations in mock community MiSeq data.**
Pairwise Spearman correlations between the vectors of estimated relative abundances for mock community samples amplified using all 96 PCR1 primer combinations (Dataset 1, Additional file 4: Table S3). Primer pairs are shown on the x and y-axis. The color of each cell indicates the degree of correlation, according to the color key on the right side of the Figure.

Additional File 13: Supplementary Figure 5
**Filename: Additional_File_13-Figure_S5_index_vs_chimera.bmp**
**Title: Association between chimeric sequence formation and reverse index sequence from PCR1.**
The data are the mock community samples amplified using 96 PCR1 primer combinations (Dataset 1, Additional file 4: Table S3). Chimera formation was not significantly associated with specific primer combinations, except for a slightly elevated rate for samples amplified with primer r13 (p<0.001, linear model). Each box represents the interquartile range while the whiskers represent 1.5 times the interquartile range. Points outside the whiskers represent outliers.

Additional File 14: Supplementary Figure 6
**Filename: Additional_File_14-Figure_S6_PCR2_PCA.bmp**
**Title: Scores plot based on a principal components analysis model computed from the matrix of species relative abundances in the mock community (Dataset 2, Additional file 5: Table S4).**
Samples are colored according to the reverse primer used for PCR1. Symbol characters represent the different PCR2 reverse primers. The first two dimensions, explaining 79% of the total variance, are shown.

Additional File 15: Supplementary Figure 7
**Filename: Additional_File_15-Figure_S7_high_low_input_boxplot.bmp**
**Title: Effects of input DNA amount used for PCR1 on relative abundance estimates in the mock community (Dataset 3, Additional file 6: Table S5).**
For enhanced visualization, each pair of colored bars (alternating blue or white for easier visualization) depicts the estimated relative abundances for one species. Species abundance estimates for high (H) and low (L) input template amounts are shown side-by-side as indicated in the x-axis labels (H=2.5e^7^ molecules, L=2.5e^6^ molecules). Each set of measurements results from 24 replicates. Each box represents the interquartile range while the whiskers represent 1.5 times the interquartile range. Points outside the whiskers represent outliers.

Additional File 16: Supplementary Figure 8
**Filename: Additional_File_16-Figure_S8_pcr1_boxplot.bmp**
**Title: Effects of PCR1 cycle number on relative abundance estimates in the mock community (Dataset 3, Additional file 6: Table S5).**
For enhanced visualization, each alternate triplet of colored bars (blue or white) depicts the estimated relative abundances for one species. Species abundance estimates for 25, 30 and 35 cycles are shown side-by-side as indicated in the x-axis labels. Each set of measurements results from 16 replicates. Each box represents the interquartile range while the whiskers represent 1.5 times the interquartile range. Points outside the whiskers represent outliers.

Additional File 17: Supplementary Figure 9
**Filename: Additional_File_17-Figure_S9_pcr2_boxplot.bmp**
**Title: Effects of PCR2 cycle number on relative abundance estimates in the mock community (Dataset 3, Additional file 6: Table S5).**
For enhanced visualization, each alternate pair of colored bars (blue or white) depicts the estimated relative abundances for one species. Species abundance estimates for 5 and 10 cycles are shown side-by-side as indicated in the x-axis labels. Each set of measurements results from 24 replicates. Each box represents the interquartile range while the whiskers represent 1.5 times the interquartile range. Points outside the whiskers represent outliers.

Additional File 18: Supplementary Figure 10
**Filename: Additional_File_18-Figure_S10_PCR1vsChimera.bmp**
**Title: Relationship between PCR1 cycle number and chimeric sequence formation in Dataset 3 (Additional file 6: Table S5).**
The number of PCR1 cycles is indicated on the x-axis. Black and red dots indicate samples amplified using 5 and 10 cycles for PCR2 respectively. A highly significant linear relationship (p<<0.001, linear regression model) was observed.

Additional File 19: Supplementary Figure 11
**Filename: Additional_File_19-Figure_S11_hsPCA1.bmp**
**Title: Effects of reverse PCR1 primer indexing in HiSeq data.**
Scores plot based on a principal component analysis model computed from the matrix of species relative abundances in the mock community sequenced on the HiSeq (Dataset 4, Additional file 7: Table S6). Samples are colored according to the reverse primer used for PCR1. The first two dimensions, explaining 72% of the total variance, are shown.

Additional File 20: Supplementary Figure 12
**Filename: Additional_File_20-Figure_S12_hsPCA2.bmp**
**Title: Effects of forward primer indexing in HiSeq data.**
Scores plot based on a principal component analysis model computed from the matrix of species relative abundances in the mock community sequenced on the HiSeq (Dataset 4, Additional file 7: Table S6). Samples are colored according to the forward primer used for PCR1. The first two dimensions, explaining 72% of the total variance, are shown.

Additional File 21: Supplementary Figure 13
**Filename: Additional_File_21-Figure_S13_correlations_hiseq.bmp**
**Title: Heat map of pairwise sample correlations in mock community HiSeq data.**
Pairwise Spearman correlations between the vectors of estimated relative abundances for mock community samples amplified using 24 PCR1 primer combinations and sequenced on the HiSeq (Dataset 4, Additional file 7: Table S6). Primer pairs are shown on the x and y axis. The color of each cell indicates the degree of correlation, according to the color key on the right side of the Figure.

Additional File 22: Supplementary Figure 14
**Filename: Additional_File_22-Figure_S14_sf_MDS.bmp**
**Title: Multidimensional scaling (MDS) plot showing clustering of 25 samples taken from five healthy adult volunteers (Dataset 6, Additional file 9: Table S8).**
Sample origin is indicated by color (Individual 1-5). The stress value of the MDS model was 13.2%, indicating a good fit.

